# Image-based Strain Analysis Reveals Intracellular Strain Controlled by Nucleo-Cytoskeletal Coupling

**DOI:** 10.1101/2024.04.18.590162

**Authors:** Jerry C.C. Chen, Iris Sloan, Alexandra Bermudez, David Choi, Ming-Heng Tsai, Lihua Jin, Jimmy K. Hu, Neil Y.C. Lin

## Abstract

Cells can sense and transduce mechanical forces, such as stretching, and convert these signals into diverse cell biological events. While much effort has been devoted to identifying the downstream biochemical and cellular responses, it is equally crucial to pinpoint the mechanical stimuli within a cell driving these responses. Specifically, much remains unknown about how intracellular strains are distributed and controlled during mechanical deformation. In this study, we developed a microscopy-based intracellular strain measurement technique. Utilizing the intrapopulation mechanical heterogeneity of epithelial monolayers, we observed an inverse relationship between cytoplasmic and nuclear strains. We found that this anti-correlation is abolished by the inhibition of Linker of Nucleoskeleton and Cytoskeleton (LINC) complex, suggesting that nucleo-cytoskeletal coupling controls intracellular strain distribution. We discovered a direct connection between cytoplasmic strain and stretch-induced nucleus size changes, implying that molecular events arising from cytoplasmic deformation may drive nuclear remodeling during stretching. By conducting multivariable analyses, we found that the intracellular strain can be inferred from cell morphology. Overall, our experimental platform and findings provide a foundation for unraveling the relationship between mechanotransduction pathways and upstream intracellular strain.

**SIGNIFICANCE:** Mechanical stimuli exert influence on epithelial cells, not only orchestrating embryogenesis and regeneration, but also regulating cancer progression and inflammatory conditions. Despite efforts to identify mechanically activated molecular events, understanding how deformation is distributed within cells to induce subcellular responses remains limited. Specifically, the control of subcellular strain distribution during mechanical stretch is unclear. In this study, we developed a microscopy-based method to measure subcellular strain and observed an inverse relationship between cytoplasmic and nuclear strains. Disrupting nucleo-cytoplasmic coupling abolished this relationship, suggesting its role in controlling strain distribution. Additionally, we found that cytoplasmic strain correlates with nucleus size changes during stretching, indicating cytoplasmic events influence nucleus remodeling.

## INTRODUCTION

Epithelial tissues experience mechanical forces during development (1, 2), injury (3–5), and regeneration (6), and these mechanical signals can have profound impacts on cell cycle progression (7), differentiation (8, 9), and migration (10). Several key mechanotransduction pathways have been identified to convert forces into biochemical signals in both physiological and pathological contexts (11–13). For instance, mechanical deformation activates ion channel Piezo1, leading to calcium influx and downstream signaling pathways that promote cell proliferation (14, 15). In myogenesis, stretching skeletal muscle tissue triggers focal adhesion kinase (FAK) and mitogen-activated protein kinase (MAPK) signaling pathways in myoblasts, facilitating their differentiation into mature muscle fibers (16–18). In cancer metastasis, extracellular matrix remodeling enhances Rho GTPase activities, promoting actin cytoskeleton reorganization and migration of cancer cells through confined spaces (19, 20). Understanding how mechanical forces govern cell behaviors provides valuable insights into fundamental cellular processes and disease progression, guiding the development of therapeutic strategies targeting these mechanical cues. To understand how mechanical stimuli, it is important to identify mechanically induced intracellular events and downstream cell phenotypes. As such, various cell stretching systems have been developed and utilized as a foundational tool. For instance, uniaxial stretching has been found to reduce nuclear stiffness through histone modification in epidermis progenitor cells (21). Biaxial stretching on mesenchymal stem cells has been found to enhance vascular regeneration through force-activated transcription factors (22). Cyclic stretching on primary cortical neurons has been found to orient the branch formation along with cytoskeleton (23).

While considerable progress has been made to uncover pathways for mechanotransduction and their downstream cellular responses, how strain is parsed within cells to deform organelles and subsequently induce subcellular responses remain unclear. During stretching, the tissue deformation influences the mechanosensors and mechanotransducers that reside at distinct subcellular locations (24). Therefore, the activities of these sensors and transducers are controlled by the intracellular strain within individual subcellular compartments, such as the cytoplasm and the nucleus (15, 25–27). Previous experiments have highlighted the intracellular mechanical heterogeneity, in which different organelles exhibit diverse moduli, implying highly non-uniform intracellular deformations (28). However, this differential intracellular strain distribution has been often overlooked in cell stretching experiments, where cellular deformation has been commonly treated as an experimental variable for all stretched cells. In these studies, attributing mechanically induced events to specific mechanical cues remains challenging. As a result, it remains relatively unknown how subcellular domains are differentially deformed to induce distinct molecular events that lead to cellular responses.

To experimentally characterize intracellular strain, light microscopy-based approaches have been widely employed due to their non-invasive nature and sub-*μ*m resolution(29). For example, quantification of intranuclear strain has revealed the heterogeneous stiffness within the nucleus and its correlation with DNA content(30). Induction and measurements of mitochondrial deformations have demonstrated the mechanosensitivity of mitochondria (31). The observation of inelastic deformations of focal adhesion complexes has revealed a novel amplification of force transmission (32).

Utilizing image-based strain quantification approaches, we developed an intracellular strain measurement platform combining a cell stretcher, live fluorescent imaging, and segmentation-based strain analysis. By stretching an epithelial monolayer on an elastic membrane, we measured its heterogeneous strain distribution in response to uniaxial stretching. Employing AI-based cell segmentation tools (33), we quantified the intracellular strain distribution, focusing on nuclear and cytoplasmic strains – two essential subcellular compartments. Leveraging the intra-population mechanical variation within an epithelial monolayer, we observed an anti-correlation between nuclear strain and cytoplasmic strain. We found that this intracellular strain distribution was abolished by disrupting nucleo-cytoskeletal coupling. Furthermore, we demonstrated that cytoplasmic strain is associated with nucleus shrinkage in cyclically stretched cells, indicating the importance of mechanotransduction pathways driven by cytoplasmic molecular activities. Lastly, we showed that intracellular strains can be inferred from morphological features of cells.

## MATERIALS AND METHODS

### 0.1 Cell culture

Plasma membrane-GFP Madin-Darby Canine Kidney (MDCK) cells and dominant-negative (DN) KASH-GFP MDCK cells were maintained at 37 °C with 5 % (v*/*v) CO_2_ and humidity in MEM-*α* (Fisher Scientific, 12561-056) supplemented with 10 % (v*/*v) fetal bonvine serum (FBS)(Fisher Scientific, 12662-029) and 1 % (v*/*v) Penicillin-Streptomycin (Fisher Scientific, 15140-1220). RPTEC-TERT cells were cultured at 37°C with 5 % (v/v) CO_2_ and humidity in DMEM*/*F12 (Fisher Scientific, 11330-032) supplemented with RPTEC growth kit (ATCC, 80320444), 10 % (v*/*v) FBS and 1 % (v*/*v) Penicillin-Streptomycin.

### 0.2 Stretching experiment

A 150 *μ*m-thick polydimethylsiloxane (PDMS) membrane attached on stretcher jigs was fabricated by our previously published procedure (34). Briefly, Sylgard 184 (base-curing agent ratio 10:1) was used for spin-coating at 2000 RPM speed for 2 min on a glass coverslip that was pre-coated with 10% (m*/*v) polyvinyl alcohol (PVA) at a speed of 1500 RPM for 2 min. Two parallel cell stretcher jigs were then attached to the uncured PDMS coating, and the composite was cured at 150°C for 30 minutes. After autoclaving, the composite was treated with 12.5 *μ*g*/*mL fibronectin (Sigma-Aldrich, F1141-2MG), and then incubated at 37°C for 30 min. The cells were then seeded on the fibronectin-treated PDMS membrane at a density of 100,000 cells per cm2 72 hours before stretching experiments to allow cell-cell junctions to mature. We then mounted and aligned the cell monolayer-jig composite on the uniaxial stretcher above the optical window, and pipetted 50 *μ*L corresponding culture media on the cell monolayer. We applied a 25 % strain single stretch or a 25 % strain 0.05 Hz cyclic stretch by controlling the stretcher motors with our customized LabVIEW program.

### 0.3 Microscopy and image analysis

Fluorescent images were acquired using a confocal microscope (RCM1 with Nikon Eclipse Ti-E, NIS-Elements software) with 20×*/*0.75 NA, 60x WI*/*1.00 NA objectives. The imaging conditions were consistently maintained across all experiments. To identify cell and nucleus contours, Z-projected images obtained from z-stack imaging using a step size of *∼*2 *μ*m were used. To reconstruct 3D nucleus volume, a fine step size *∼*0.2 *μ*m were used for z-stack imaging. To measure the strains and morphology, the projected images at different stretching states were registered to remove offset by the registration tool in Fiji. Then, cell and nucleus segmentation was processed by Cellpose 2.0(33) to generate the respective object masks. The masks of individual cells and nucleus were paired and labeled by the customized Macros code in Fiji. Lastly, nuclear *xx* strain (*ϵ*_*n*_), cytoplasmic *xx* strain (*ϵ*_*c*_), cellular *xx* strain (*ϵ*_*cell*_) (Δ*u/*Δ*x*) across the nuclear centroid and the morphological features were measured by our customized Matlab code.

### 0.4 MDCK cell transductions and inhibition experiments

To tag the plasma membrane with GFP, wild-type (WT) MDCK cells were transfected with pAcGFP1-Mem vector (Takara, 632509). To decouple the LINC complex by overexpressing dominant-negative KASH protein, WT MDCK cells were transfected with GFP-KASH2 vector (Addgene, plasmid no. 187017) using Lipofectamine Reagents (Invitrogen, 18324012) and PiggyBac transposon vector. After neomycin selection, FACS (BD FACS Aria H) was used to sort the positive clonal populations.

To inhibit myosin II and microtubules, the cells were treated with blebbistatin (Millipore Sigma, B0560) at 25 *μ*M for 2 hr and nocodazole (Millipore Sigma, M1404) at 10 nM for 24 hr before the mechanical stretching, respectively. These optimal concentrations were determined by titrating doses to achieve their primary functions without causing other adverse effects. The DN-KASH MDCK line was treated with doxycycline at 1 *μ*g*/*mL to induce the overexpression of DN-KASH.

### 0.5 Canonical correlation analysis

The canonical correlation analysis (CCA) was conducted using a Matlab built-in function (canoncorr). The deformation canonical variate was a linear combination of the three strains given by the respective weights. Similarly, the morphology canonical variate was a linear combination of the nineteen morphological features given by the respective weights. By calculating the maximum correlation between deformation canonical variable and morphology canonical variable, the weights were determined and the relationship between factors were examined. Three correlation sets were generated due to the minimum number of the factors, which were the three strains.

### 0.6 Data analysis and statistics

Statistical analyses were performed using GraphPad PRISM (version 10). For correlative analyses of all presented scatter plots, the Pearson correlation coefficient, confidence interval (CI) and the p-value were calculated. To examine the relationship between intracellular strains, False discovery rate (FDR) was evaluated using a Monte Carlo simulation in Matlab that randomly permutes the X Y values of data points. P-values > 0.05 are denoted as not significant (ns), while p-values *≤* 0.05, *≤* 0.01, *≤* and *≤* 0.0001, are represented as *, **, ***, and ****, respectively.

## RESULTS

### Live fluorescent imaging of cell stretching enables intracellular strain measurements in epithelial monolayers

In order to precisely measure how cells deform in response to stretching, we developed a subcellular strain deformation measurement platform that integrates cell stretching, live fluorescent imaging, and segmentation-based image analysis. Our custom-designed cell stretcher, featuring dual motors on opposite sides, applies symmetric uniaxial deformation with encoder-based closed-loop control over stretching rate and amplitude (Fig. S1a) (34). To conduct the strain measurement, we first grew a confluent monolayer of membrane-GFP Madin-Darby canine kidney (MDCK) cells on a 150*μ*m-thick polydimethylsiloxane (PDMS) membrane, which was then affixed to the cell stretcher and mounted on an inverted fluorescent microscope (Fig. 1a). The sample was stained with Hoechst to label nuclei 30 minutes before stretching. Beneath the mounted sample, we engineered an imaging window for live fluorescent imaging during stretching. Throughout this work, we applied a step-wise *∼*25% tensile strain, in which the actual deformation was then characterized using the acquired images (Fig. S1b).

**Figure 1:**
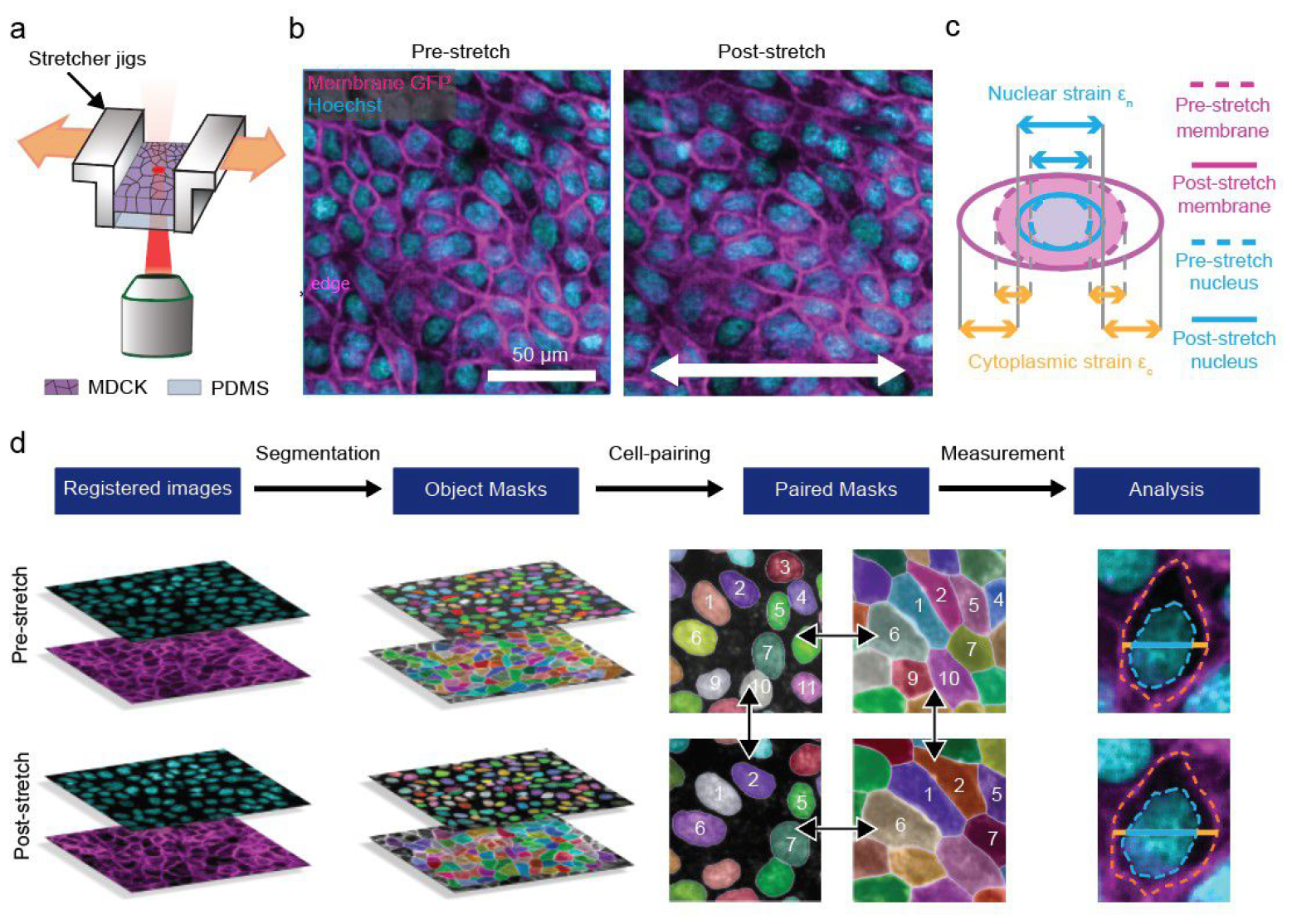
Microscopy-based intracellular strain measurements in epithelial monolayers. (a) An MDCK monolayer was cultured on a PDMS membrane. The MDCK monolayer-PDMS composite was then mounted on a custom-built cell stretcher which applied a uniaxial tensile strain while allowing simultaneous confocal imaging. (b) Live fluorescent images of the pre-stretch (left) and post-stretch (*∼*25% strain, right) MDCK monolayer. Stretched cells exhibited nuclear and cellular morphological elongation along the axis of the applied strain. (c) Schematic illustrating the definitions of nuclear and cytoplasmic strain along the stretching direction used in this work. (d) Workflow overview outlining strain analysis pipeline. Cell and nucleus images were first registered before segmentation to obtain the respective object masks. Within the same condition (e.g., pre-stretch or post-stretch), cells were then paired with their corresponding nucleus prior to matching pre-stretch cells to the same cell in the post-stretch image. Lastly, cell and nucleus strains were quantified as illustrated in (c).

To characterize the intracellular strain distribution, we focused on quantifying the deformation of nucleus and cytoplasm, two major cellular compartments regulating distinct mechanically induced molecular events (15, 25–27). As shown in Fig. 1b, by capturing fluorescent images of membrane-GFP and Hoechst, we visualized shape changes of cells and nuclei due to the applied uniaxial deformation along the horizontal direction (i.e., x-axis). While the fluorescent image pair allows the determination of the complete planar strain tensor, we focused on the *xx* strain since it is the dominant component in our experiment. Here, we defined the nuclear *xx* strain, *ϵ*_*n*_, as the ratio of nucleus length change to its initial length, and the cytoplasmic *xx* strain, *ϵ*_*c*_, as the ratio of cytoplasm length change to initial length (Fig. 1c). We summarized our image-based strain measurement workflow in Fig. 1d. In brief, all obtained images were first registered to remove offset between image acquisitions. Using a recently developed cell segmentation tool, Cellpose (33), we generated object masks for individual nuclei and cells to identify their contours. These masks were then paired between fluorescent channels and acquisitions, allowing integrated morphological analyses. The extracted contours were then used to quantify corresponding strain values and morphological features.

Image analyses reveal an

### Anti-correlation between nuclear strain and cytoplasmic strain

To investigate the relationship between the cytoplasmic and nuclear strains in response to the tensile stretch, we plotted the cellular, nuclear, and cytoplasmic strains for 300 cells in Fig. 2a. We found that the mean cellular strain was approximately 24%, consistent with the applied 25% strain. The mean nuclear strain was roughly half of the mean cellular strain, while the mean cytoplasmic strain was roughly two times of the mean cellular strain, consistent with previous stiffness measurements showing that the nucleus is stiffer than the cytoplasm(35, 36). By showing the normalized histograms of cellular, nuclear, and cytoplasmic strains, we found that all three strain components exhibit a broad unimodal distribution (Fig. 2b). To investigate the distribution of these intracellular strains, we plotted the cytoplasmic strain versus nuclear strain for all analyzed cells. As shown in Fig. 2c, this scatter plot revealed that the heterogeneous intracellular strains were not merely random but showed a moderate anti-correlation with a Pearson’s correlation coefficient -0.32, p-value < 0.001, and FDR = 0 (Fig. S2). Our observed anti-correlation implies that cells with large nuclear strains tend to exhibit smaller cytoplasmic strains, and vice versa. We further confirmed the statistical significance of this finding by comparing the cytoplasmic strains between cells that exhibited small (0-15%) and large (30-50%) nuclear strains, in which the p-value was <0.001 (Fig. 2d). We made the same observation of anti-correlation using an immortalized proximal tubule epithelial cell line (RPTEC-TERT), as shown in Fig. 2e, suggesting that this phenomenon is not unique to MDCK cells.

**Figure 2:**
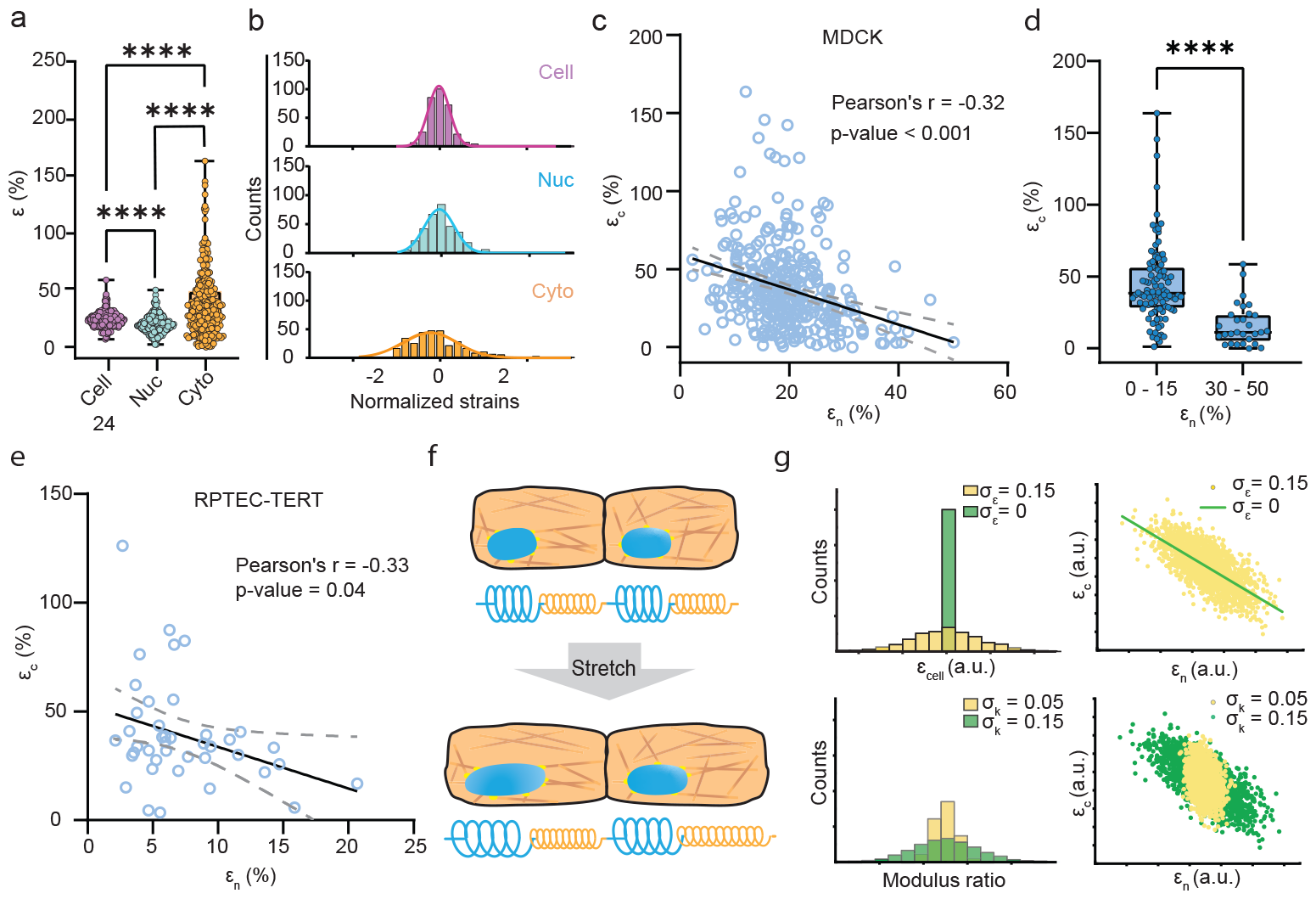
Anti-correlation between nuclear strain and cytoplasmic strain. (a) Cellular, nuclear, and cytoplasmic strains for MDCK cells stretched 24%. The mean cellular strain *∼*24% was consistent with the applied tensile strain, while the mean nucleus and cytoplasmic strains were respectively two times lower and higher than the applied strain. N > 300 cells. (b) Normalized histograms for cellular, nuclear, and cytoplasmic strains exhibited unimodal distributions with varying degrees of heterogeneity. Each strain was normalized to its mean. Solid lines denote a Gaussian fit for each histogram. (c) Scatter plot of nuclear strain vs. cytoplasmic strain shown in (a) and (b) indicated an anti-correlation between the two strains. Solid line and dotted lines denote the best fit and 95% confidence interval, respectively. (d) Anti-correlation between the nuclear and cytoplasmic strains was confirmed by the statistically significant difference in cytoplasmic strain between the low (0-15%) and high (30-50%) nuclear strain groups. (e) RPTEC-TERT cells exhibited a qualitatively similar nuclear and cytoplasmic strain anti-correlation. N > 40 cells (f) Schematics of nucleus and cytoplasm transmitting force in series under a global deformation. Blue and orange springs represent the nuclear and cytoplasmic compartments, respectively. (g) Monte Carlo simulations demonstrating that nucleus-cytoplasm force balance partitions intracellular strain. As illustrated in the upper two plots, a system with a constant cellular strain (*σ*_*ϵ*_ = 0, green) exhibits a perfect nucleus-cytoplasm strain anti-correlation while heterogeneity in cellular strain (e.g., *σ*_*ϵ*_ = 0.15, yellow) attenuates the magnitude of the correlation. As illustrated in the bottom two plots, heterogeneity in the nucleus-to-cytoplasm stiffness ratios (*σ*_*k*_) promotes a robust anti-correlation. Here, *σ*_*ϵ*_ and *σ*_*k*_ denote the standard deviations of cellular strain and nucleus-to-cytoplasmic modulus ratio, respectively. The simulated strain has an arbitrary unit (a.u.).

Our results above indicate that the intracellular strain distribution arises from a force balance between nucleus and cytoplasm, a notion that is consistent with previous studies (37–39). To explore this phenomenon, we employed a one-dimensional spring system, where nucleus and cytoplasm are considered as two springs in series (Fig. 2f). For simplicity, we did not consider any cellular mechanical properties such as viscoelasticity, plasticity, Poisson’s ratio, or mechanical remodeling during stretch (40–42). We first considered an extreme case, where all cells exhibit identical strains (*σ*_*ϵ*_ = 0), and used Monte Carlo simulations to randomly generate 2000 samples. In all simulation cases, we assumed that the length of the nucleus and cytoplasm was equal. The first case yielded a perfect *ϵ*_*c*_ *− ϵ*_*n*_ anti-correlation (Fig. 2g, upper row, green line). Such a result is anticipated, as *ϵ*_*c*_ and *ϵ*_*n*_ add to a constant when assuming equal lengths for the nucleus spring and the cytoplasm spring. To implement a condition that better reflects cellular mechanical heterogeneity, we then considered a case, where cellular strains obey a Gaussian distribution with a standard deviation *σ*_*ϵ*_ = 15%. We found that the anti-correlation is weakened but persisted (Fig. 2g, upper row, yellow dots). We further examined how intracellular mechanical heterogeneity affects intracellular strain distribution by varying the range of nucleus-to-cytoplasm strain ratio (i.e., stiffness ratio). We found that a uniform ratio (i.e., narrow range) resulted in a weak anti-correlation (Fig. 2g, lower row, yellow dots). In contrast, a broad distribution of nucleus-to-cytoplasm strain ratio gave rise to a robust anti-correlation. In summary, our simulations suggested three essential conditions underlying the *ϵ*_*c*_ *− ϵ*_*n*_ anti-correlation: (1) mechanical coupling between nucleus and cytoplasm, (2) unimodal distribution of cellular stiffness, and (3) intracellular mechanical heterogeneity.

### Nucleo-cytoskeletal coupling controls intracellular strain distribution

As suggested by our simulation, the strain partitioning between the nucleus and the cytoplasm arises from their mechanical coupling. To examine this intracellular strain distribution mechanism, we disrupted three intracellular force transmission components, actomyosin, microtubules, and LINC, as summarized in Fig. 3a. First, we weakened actin tension by administering Blebbistatin (Bleb) to the sample, reducing the binding of non-muscle myosin II to actin filaments(43). We mitigated microtubule formation using Nocodazole (Noco), which inhibits the polymerization of microtubules(44). Lastly, we disrupted nucleo-cytoskeletal coupling by overexpressing a dominant-negative KASH-GFP (DN-KASH) construct (45, 46), which blocks binding of endogenous SUN and Nesprin proteins of the LINC complex that physically connects the nuclear interior to the cytoskeleton. We first examined how these inhibitions affected intracellular force transmission by measuring the mean normalized nuclear strain. Our measurements show that both Blebbistatin and DN-KASH significantly reduced nuclear strain compared to control, while Nocodazole showed no significant effect (Fig. 3b). The effect of Bleb was further validated by morphological and atomic force microscopy measurements, in which Bleb reduced the cell modulus and altered cell morphology (Fig. S3).

**Figure 3:**
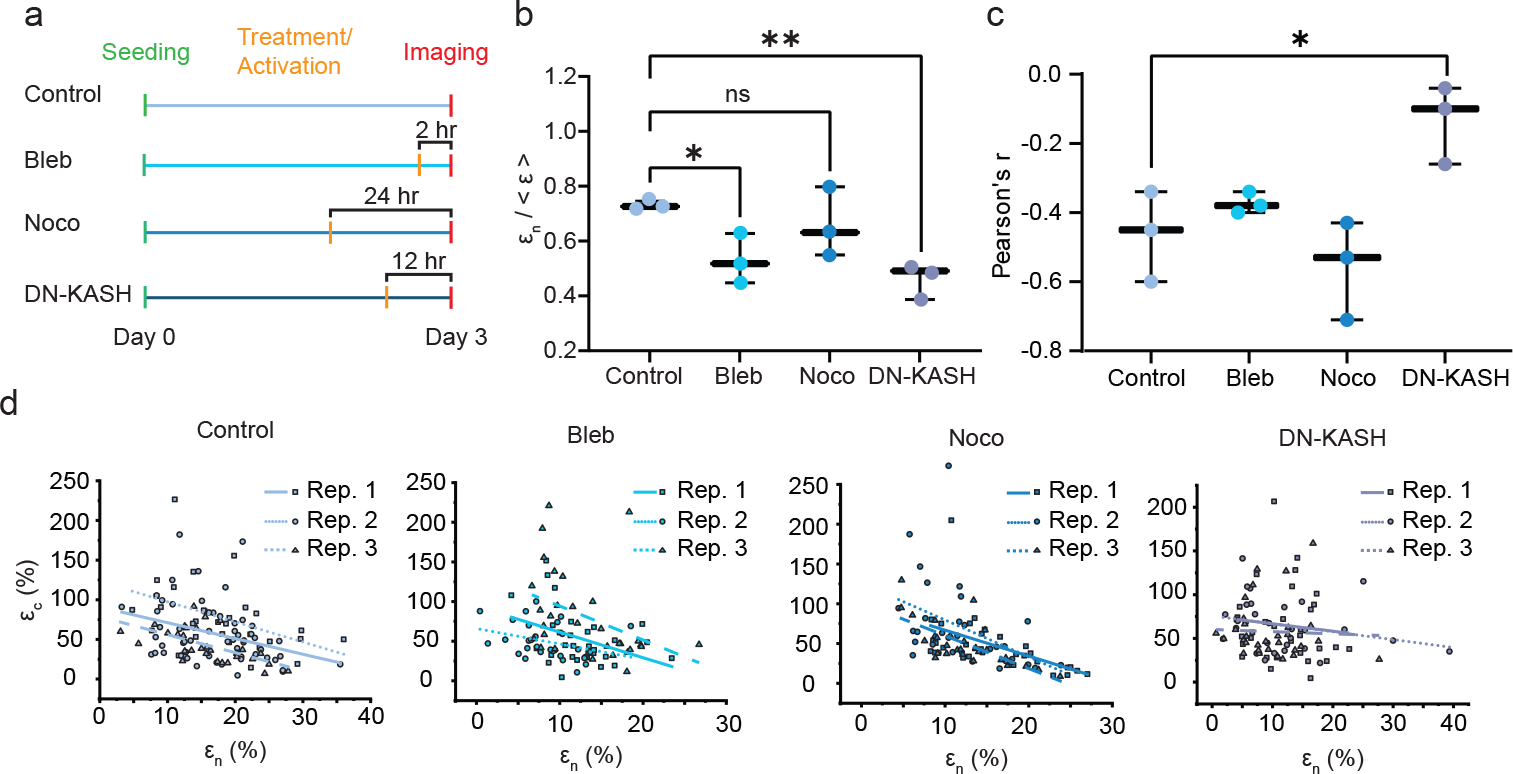
Nucleo-cytoskeletal coupling is required for intracellular strain distribution control. (a) Experiment overview investigating the role of cytoplasmic force propagation in the anti-correlation between nuclear and cytoplasmic strain. Blebbistatin (Bleb), nocodazole (Noco), and a dominant-negative KASH GFP reporter line (DN-KASH) were used for Myosin-II inhibition, microtubuele polymerization inhibition, and LINC complex disruption respectively. (b) Relaxation of actomyosin tension and disruption of LINC complex reduce the normalized nuclear strain while microtubeles polymerization inhibition play a secondary role in regualtig the nucelar strain. Nuclear strains were normalized to the global tensile strain of each condition. N = 3. (c) The physical decoupling of the nucleoskelton to the cytoskeleton reduces the magnitude of the anti-correlation between nuclear and cytoplasmic strain while the relaxation of the cytoskeleton does not play a role in generating the observed anti-correlation. Scatter plots illustrating the anti-correlation between cytoplasmic strain and nuclear strain for (d) Control (e) Bleb (f) Noco (g) KASH groups. A physical connection between the nucleoskeleton and the cytoskeleton regualted the cytoplasmic strain and nuclear strain correlation. Solid lines represent the best fit line.

To investigate how inhibitions of actomyosin, microtubule, and nucleo-cytoskeletal coupling interfere with the intracellular strain distribution, we analyzed the nucleus-cytoplasm strain correlation, and summarized the Pearson correlation coefficient summary in Fig. 3c. We also showed the individual scatter plots for all conditions in Fig. 3d. Representative images of the control and stretched samples are shown in Fig. S4. Our correlation analysis found that the anti-correlation was undisrupted in the Bleb and Noco samples, suggesting that actomyosin tension and microtubule assembly are not required for intracellular strain correlation (29, 47–49). In contrast, the anti-correlation was significantly attenuated in the DN-KASH group, as indicated by its near-zero Pearson correlation coefficient. This finding indicates that nucleo-cytoskeletal coupling is required for intracellular strain distribution control, consistent with previous findings that LINC complex plays a crucial role in force transmission to the nucleus and governs nuclear mechanosensing (38, 50, 51).

### Nuclear and cytoplasmic strains are deferentially associated with nuclear size change

Nuclear size change has been shown to be a prominent cellular phenotype alteration in response to mechanical stretch (52–54). We thus sought to understand how intracellular strain distribution affects nuclear size change, since force-activated molecular events in different subcellular compartments can be distinct. Following published experiments (55, 56), we performed cyclic stretching to recurrently deform an epithelial monolayer (Fig. 4a), to be consistent with their protocols. In this experiment, we first characterized the nuclear and cytoplasmic strains across one field of view during the first stretch cycle. We then conducted 100 stretch cycles and subsequently acquired fluorescent images for nuclear size measurements. To approximate nuclear size, we used the projected area, validating this approach by observing excellent agreement between 3D nuclear volume and projected area in a confocal image stack (Fig. S5). While initial analysis showed no significant difference in mean nuclear area between pre-stretch and cyclically stretched samples (Fig. 4b), we scrutinized individual nuclei and found that the nucleus could either shrink (i.e., *≥*1% area decrease), maintain its size (i.e., <1% area change), or expand (i.e., *≥*1% area increase) in response to cyclic stretching (Fig. 4c). This diverse nuclear response is illustrated by the broad distribution of the area change histogram (Fig. 4d). This observation of both shrinking and expanding nuclei is consistent with previous findings that mechanical stretch induces various molecular responses that can lead to either an increase or a reduction in nuclear size (21, 56, 57).

**Figure 4:**
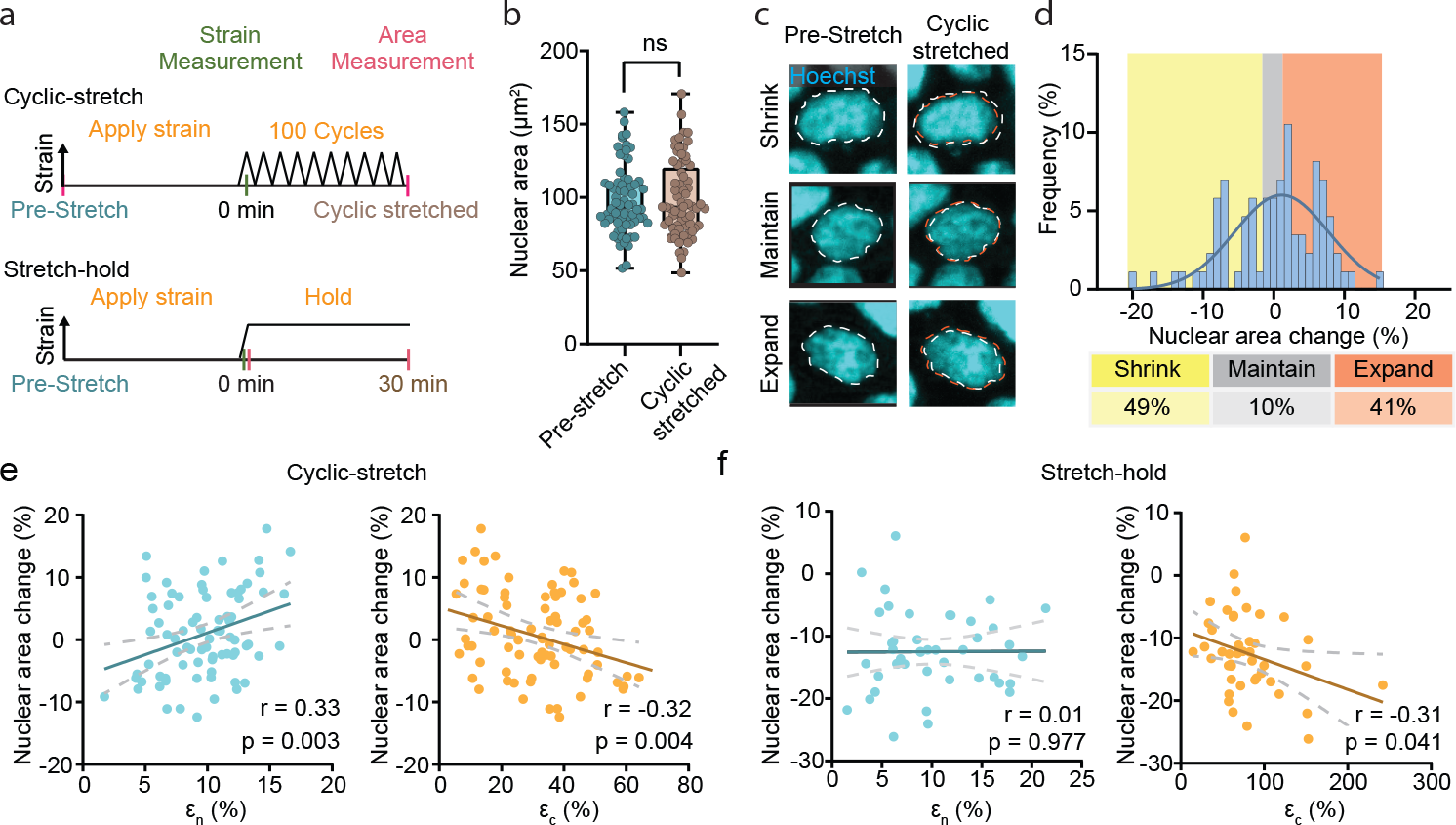
Stretch-induced nuclear size change is regulated by intracellular strain distribution. (a) Experiment overview of two different stretching profiles applied to MDCK cells. In one experiment, the initial nuclear area was measured before applying 25% strain to the monolayer and then measuring the resulting strain. Following this, the monolayer was cyclically stretched 100 times at a frequency of 0.05 Hz prior to measuring the nuclear area. In a separate experiment, 25% strain was applied to the monolayer and the resulting strain and nuclear area was measured. The sample was then held in the stretched state for 30 minutes and the nuclear area was remeasured. (b) Nuclear area change in response to cyclic stretching shows no significant difference. (c) Fluorescent images of nuclei before and after cyclic stretching. Cyclic stretching induces a heterogeneous response in nuclear size change as shrinking, no change, and expansion we observed. (d) Histogram illustrating nuclear size response to cyclic stretching. Gaussian fit overlaid. (e) Scatter plots of nuclear area change versus nuclear strain *ϵ*_*n*_ (left) and nuclear area change versus cytoplasmic strain *ϵ*_*c*_ (right) respectively demonstrate a positive and negative correlation when cyclically stretched. (f) Scatter plots of nuclear area change versus nuclear strain *ϵ*_*n*_ (left) and nuclear area change versus cytoplasmic strain *ϵ*_*c*_ (right), respectively.

To investigate how nuclear size changes may be governed by intracellular strain, we plotted the area change versus nuclear strain (*ϵ*_*n*_) or cytoplasmic strain (*ϵ*_*c*_) as shown in Fig. 4e. We found that nuclear area change was positively correlated to nuclear strain (Pearson’s correlation coefficient *∼* 0.33), but negatively correlated to cytoplasmic strain (Pearson’s correlation coefficient *∼* -0.32). The observed correlations indicate that intracellular strain distribution influences nuclear size change during stretching. This finding also put forth a hypothesis for future studies, in which we posit that nuclear expansion is mainly driven by nuclear deformation, while nuclear shrinkage is mainly caused by cytoplasmic deformation.

We also preformed a stretch-hold experiment (Fig. 4a) and repeated the nuclear area analysis, since stretch-hold represents a different way to mechanically perturb cells and can elicit different mechanical and cellular responses. We found that while the stretch-hold result (Fig. 4f) was qualitatively consistent with the cyclic-stretch results (Fig. 4e), most nuclei exhibited an area reduction in the stretch-hold experiment as shown in Fig. S6a (The representative images of nuclear area change are in Fig. S6b). Consequently, the correlation between nuclear area change and *ϵ*_*n*_ was low in the stretch-hold experiment, consistent with our findings in the cylic stretch experiment (replication results shown in Fig. S6c, f). Collectively, we showed that nuclear and cytoplasmic strains play differential roles in regulating nuclear mechanoresponse.

### Intracellular strain response can be inferred from cell morphology

Thus far, we have shown that the intracellular strain within each cell is closely regulated by nucleo-cytoskeletal coupling and is important for nuclear mechanoresponses. We next asked whether such strain distribution can be predicted by cell morphology across a stretched monolayer. Similar to the strain response, cell morphology is a phenotype directly governed by cell mechanical properties, such as nucleus stiffness, cortical tension, and cytoskeletal viscoelasticity(58–60). Therefore, we conjectured that the morphological features of an unstretched cell layer provide sufficient information to infer the strain distribution in its stretched state. To this end, we investigated the association between cell deformation and morphological features using Canonical Correlation Analysis (CCA)(61). As illustrated in Fig. 5a, CCA identified the linear combinations of variables from the deformation canonical variables (v) and morphology canonical variables (u) that maximize the correlation between them, revealing the shared variation between the two sets. Here, the deformation canonical variable set contains *ϵ*_*n*_, *ϵ*_*c*_, and *ϵ*_*cell*_ and the morphology canonical variable set contains 19 morphological features of cell and nucleus, where representative examples are shown in Fig. 5b and Table S1. Furthermore, since all cells are physically connected in a confluent layer, their strain responses depend on the mechanical properties of all cells. We thus conducted an additional CCA that considers such a multicellular effect. For simplicity, we included the morphological features of the three nearest neighboring cells to account for intercellular mechanical interactions (Fig. 5c). Each CCA result reports three correlation coefficients.

**Figure 5:**
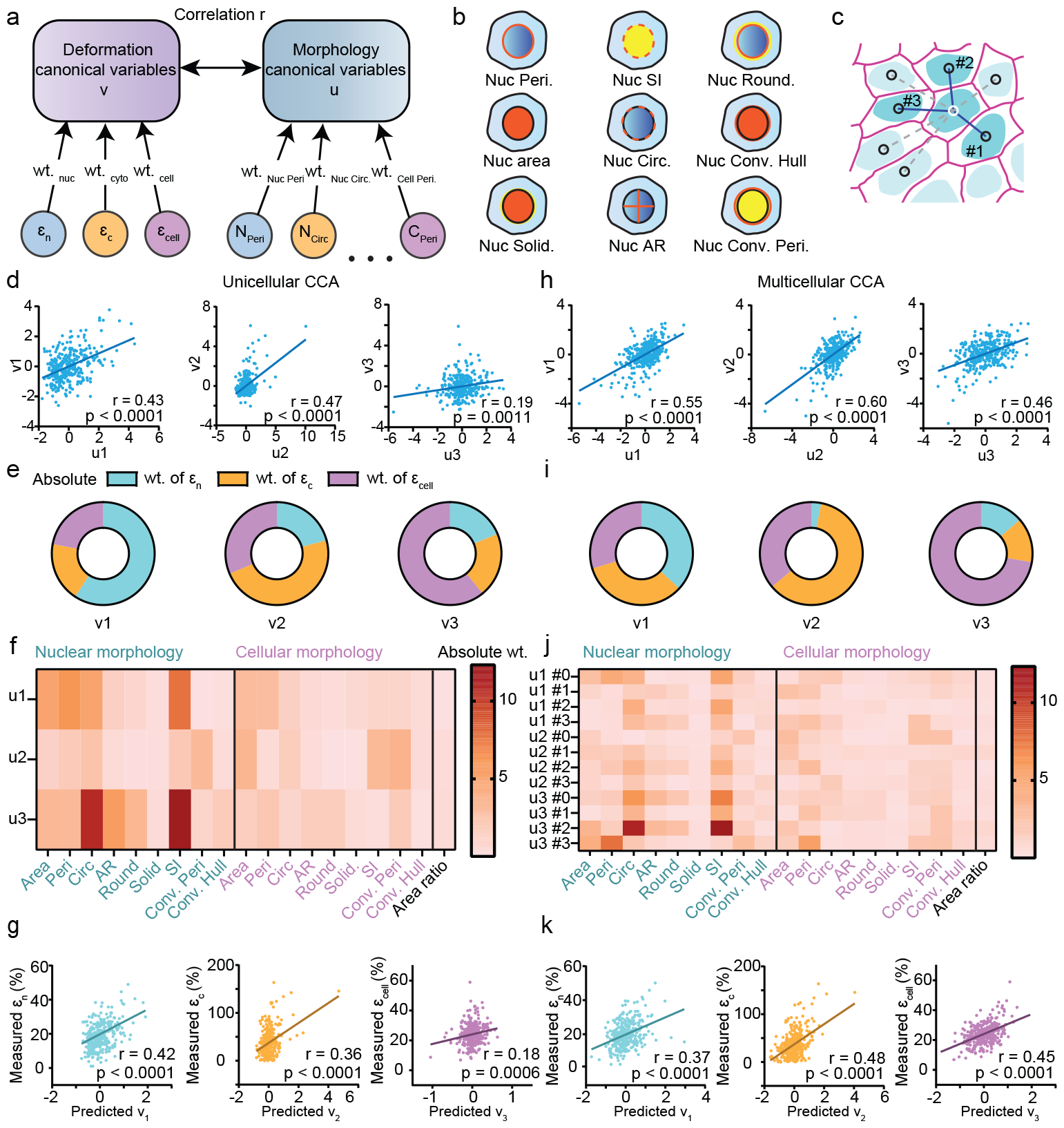
Strain distributions are correlated with cell morphology. (a) Illustration of deformation (v) and morphology (u) variables used for the canonical correlation analysis (CCA), which determines the maximum correlation between u and v by optimizing the weights (wt.) of the canonical variables. (b) Schematics of representative morphology canonical variables, including perimeter (Peri.), area, solidity (Solid.), shape index (SI), circularity (Circ.), aspect ratio (AR), roundness (Round.), convex hull (Conv. Hull) and convex perimeter (Conv. Peri.). (c) Schematic illustrating that the top 3 closest neighbor cells (#1-3) were included in the multicellular CCA. (d) Scatter plots of canonical variates for unicellualr CCA. (e) The relative weights of nuclear strain, cytoplasmic strain and cellular strain for v1, v2 and v3. (f) Heatmap of the weights of the morphological features for u1, u2 and u3. (g) Scatter plots of measured strains vs. predicted variates for unicellular CCA. (h) Scatter plots of canonical variates for multicellualr CCA, which exhibited higher correlations compared to the unicellular approach. (i) The relative weights of nuclear strain, cytoplasmic strain and cellular strain for v1, v2 and v3. (j) Heatmap of the weights of the morphological features for u1, u2 and u3, as well as the target cell (#0), closest (#1), second-closest (#2), and third-closest (#3) neighbor cells. (k) Scatter plots of measured strains vs. predicted variates for multicellular CCA

In the unicellular CCA, we found that the v-u correlations are higher for the first and second combinations, or variates (Fig. 5d) with nuclear strain weight or cytoplasmic strain weight dominating over the other two deformation variables (Fig. 5e). The third variate showed a relatively weak correlation where cellular strain carried dominant weight. In analyzing the weight distribution in the morphology variable (u) set, we found that nuclear morphological features were consistently given heavier weights than cell morphology, indicating that the nuclear morphological features such as circularity (area/perimeter2) and shape index (i.e., perimeter/area1*/*2) are essential for inferring intracellular deformation (Fig. 5f). To confirm the predictability of nuclear strain and cytoplasmic strain as shown in Fig. 5e, we plotted the measured strain against CCA-predicted variate value (Fig. 5g), and found that nuclear strain *ϵ*_*n*_ is in good agreement with v1 (Pearson correlation coefficient *∼* 0.42), whereas cytoplasmic strain *ϵ*_*c*_ is in good agreement with v2 (Pearson correlation coefficient *∼* 0.36). However, the total cell strain *ϵ*_*cell*_ was not accurately predicted.

In the multicellular CCA, we observed an overall improvement in correlations between deformation and morphology (Fig. 5h). Compared to the unicellular analysis, both cytoplasmic strain and cellular strain were more weighted in multicellular CCA (Fig. 5i). Consistent with the unicellular CCA, we still found that nuclear morphology features carried more weights than cellular features (Fig. 5j). Importantly, we observed approximately equal weights between the cell of interest (#0) and its neighbors (#1, #2, and #3), confirming the importance of including neighboring cells (62). Lastly, we showed that cytoplasmic strain and cellular strain were more accurately predicted by morphology in multicellular CCA, showing Pearson correlations coefficient *∼* 0.48 and 0.45, respectively (Fig. 5j). Altogether, our CCA results indicate that the nuclear strain of a stretched cell can be predicted by its own morphology, whereas accurate inferences of cytoplasmic strain and cellular strain require the morphological information of surrounding cells.

## CONCLUSION AND DISCUSSIONS

In this study, we introduced an experimental approach for measuring intracellular strain in stretched epithelial monolayers. Using our intracellular strain measurements, we demonstrated the biomechanical role of nucleo-cytoskeletal coupling in governing intracellular strain distribution. Prior studies have revealed the nucleo-cytoskeletal coupling’s function in transmitting extracellular force to the nucleus (38, 50, 51), thereby inducing nuclear remodeling (63), chromatin modification (21), and alterations of gene expression (64). Our work sheds light on its additional function in modulating the ratio of nuclear to cytoplasmic strains. We primarily focused on investigating the LINC complex by disrupting Nesprin-SUN binding. The connection between the nucleus and cytoskeleton along with the intracellular force balance collectively result in our observed anti-correlation between nuclear and cytoplasmic strains. In future experiments, it would be valuable to examine other nucleo-cytoskeletal coupling pathways. For instance, the actin cap over the nucleus can physically restrict the nucleus’s vertical dimension, establishing a mechanical coupling between the nucleus and cytoplasm (65, 66). Additionally, the endoplasmic reticulum, which merges with the nucleus membrane and spatially aligns with the nucleus, can serve as a connection between the cell membrane and nucleus (67).

To consider all potential mechanisms responsible for the anti-correlation reduction besides the nucelo-cytoskeletal coupling disruption, we noted that the overall nuclear strain was reduced in both Bleb and DN-KASH groups, potentially due to the attenuated force-transfer rate. Such a reduced nuclear strain then led to a narrowed distribution of strain ratio (i.e., stiffness ratio), which might weaken the anti-correlation, as suggested by our Monte Carlo simulations. This contribution could potentially explain the slight reduction in the anti-correlation upon myosin II inhibition (Fig 3c). However, since the nucleo-cytoskeletal decoupling will always lead to reduced nuclear strain and altered strain ratio distribution, we envision that deciphering these effects will require a combination of theoretical modeling and experiments (48, 68), in which the nucleo-cytoskeletal coupling is systematically perturbed and the resulting intracellular distribution is measured.

Our study also elucidated the distinct links between nuclear strain, cytoplasmic strain, and stretch-induced nucleus size change, a phenomenon conventionally hypothesized to be a nuclear mechanosensing response safeguarding DNA from mechanical damage (54, 56). Our finding suggests that nuclear strain and cytoplasmic strain may trigger molecular events that differentially regulate nuclear size, paving ways for future mechanistic studies. For example, nuclear strain might stretch the nuclear pore complex, known to facilitate the importation of YAP and chromatin-modifying enzymes, potentially augmenting nuclear size through increased intranuclear osmotic pressure (69, 70). Furthermore, nucleus deformation has been observed to untether peripheral chromatin, possibly leading to increased entropic pressure (21, 71). In parallel, cytoplasmic strain is associated with focal adhesion activation, which promotes actin polymerization, potentially deforming the nucleus through actin cap formation (56, 72). Moreover, mechanical stimuli on the endoplasmic reticulum have been shown to trigger calcium influx, which can reduce nuclear size by increasing cytoplasmic osmotic pressure or inducing chromatin condensation (15, 73–75). These hypotheses can be explored by integrating our intracellular strain measurements with pharmacological inhibitions (76), osmotic shock techniques (77), and ultra-fine structure microscopy (78).

We further demonstrated that cell morphology features can be used for inferring the intracellular strain response. This finding aligns with recent experiments illustrating a strong connection between cell morphology and response to mechanical (34) and biochemical (79) stimuli. Our results also suggested that cell-cell mechanical connection regulates individual cell deformation in a stretched confluent epithelium, as evidenced by the enhanced prediction accuracy in multicellular CCA. This observation is consistent with previous measurements of supracellular strain (34), emphasizing the significance of multicellular mechanical coupling.

We acknowledge that both stress and modulus distributions remain unknown in our experiment, since they are independent to the strain field, and at least one of them must be measured to determine the other. While it has been suggested that stress measurement could provide more relevant information for mechanotransduction when cells are stiff and do not exhibit a significant strain response (80), we find strain measurement is still informative as cellular deformation is primarily governed by the applied tensile strain. Moreover, measuring nuclear and cytoplasmic strains estimate the deformation and subsequent activation of mechanosensors and mechanotransducers, since studies have shown that they have relatively minor impacts on the stiffness of their surroundings (15, 25, 69, 81). For instance, the cytoplasmic modulus primarily arises from cytoskeletal components(48), while nuclear stiffness is mainly influenced by properties of the nuclear envelope and chromatin packing(60). In future experiments, integrating molecular strain sensors, such as fluorescence resonance energy transfer (FRET) probes (82) and optical membrane tension sensors (83) tagged to specific mechanosensors or mechanotransducers in our system could provide valuable insights into molecular-level strain dynamics.

Overall, our work introduced an innovative image-based strain measurement and revealed the nucleo-cytoskeletal control of intracellular strain, which controls the nuclear size change in stretched monolayers. By leveraging the diverse morphological phenotype and intracellular deformations, we employed multivariable analyses to establish a link between cell morphology and mechanical response. We envision that intracellular strain measurements will be essential for deciphering the complex interplay between upstream mechanical stimuli and downstream mechanotransduction pathways.

## AUTHOR CONTRIBUTIONS

JCCC, AB, JKH, and NYCL designed the research. JCCC, IS, DC, and MHT carried out experiments. JCCC, IS, AB, LJ, JKH, and NYCL analyzed the data. JCCC, IS, AB, LJ, JKH, and NYCL wrote the article.

## ACKNOWLEDGMENTS

NYCL was supported by the UCLA California NanoSystems Institute and the Noble Family Innovation Fund, the National Science Foundation (NSF CMMI-2029454, CBET-2244760, and DBI 2325121), and the National Institute of Health NIGMS (1R35GM146735). LJ acknowledges a National Science Foundation MPS-NCI SPARK supplement (No. 2326800) to her CAREER Award (No. 2048219). J.K.H. was supported by NIH NIDCR (R01DE030471).

## SUPPLEMENTARY MATERIAL

An online supplement to this article can be found by visiting BJ Online at http://www.biophysj.org.

