## Supplemental Information for "Image-based Strain Analysis Reveals Intracellular Strain Controlled by Nucleo-Cytoskeletal Coupling"

#### Supplement information

### Image-based Deformation Analysis Reveals Intracellular Strain Distribution Controlled by Nucleo-Cytoskeletal Coupling in Stretched Epithelial Monolayers

**Jerry C.C. Chen, Iris Sloan, Alexandra Bermudez, David Choi, Ming-Heng Tsai, Lihua Jin, Jimmy K. Hu, and Neil Y.C. Lin**

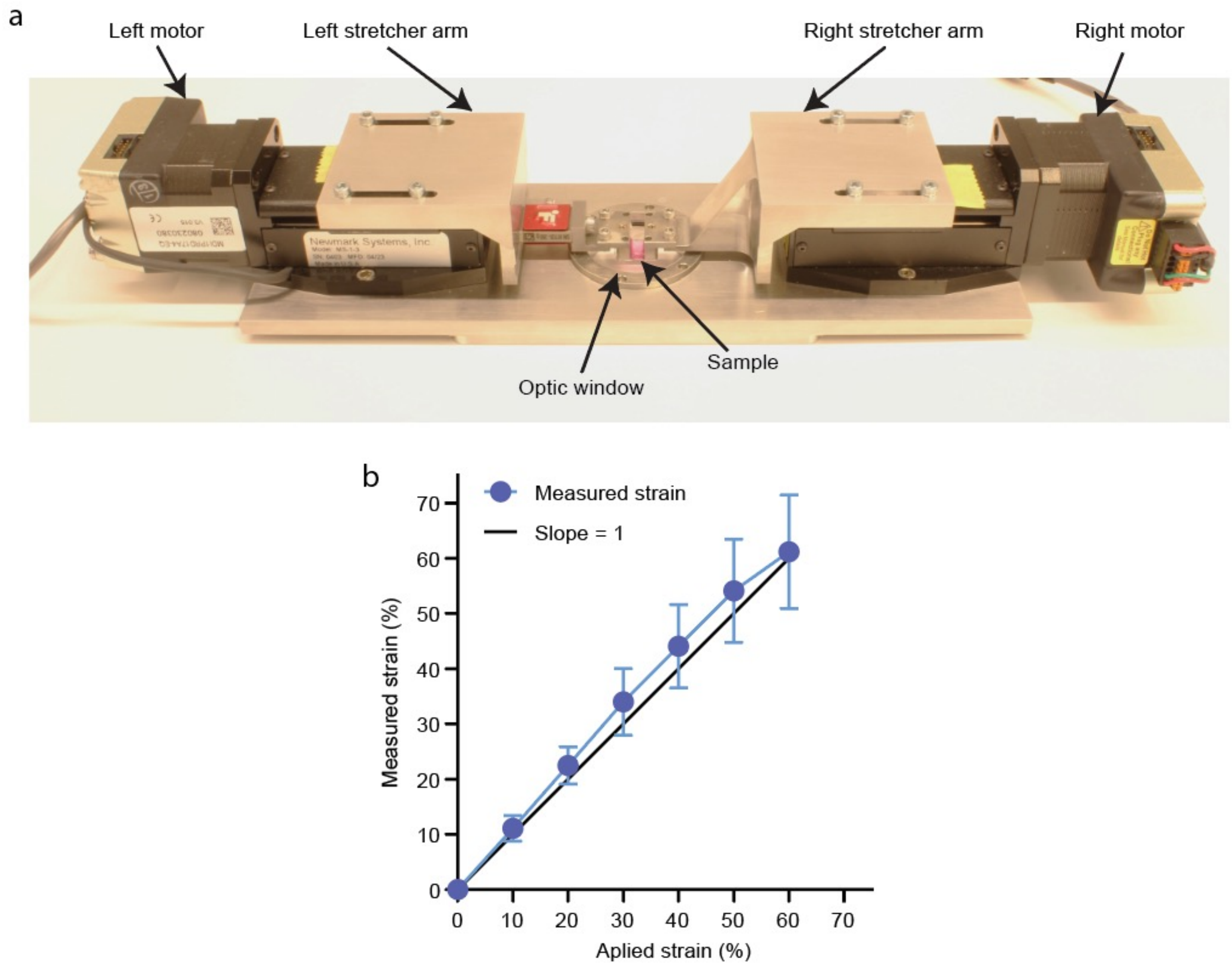


**Figure S1. Cell stretcher and strain calibration.** (a) The images of our cell stretcher with dual motors and arms to apply a symmetric uniaxial stretch on the cell monolayer-membrane composite. The optic window below the sample allows live imaging of the cell monolayer. (b) The strains measured by cell monolayer deformation at three locations in fluorescent images were consistent with the applied tensile strain.

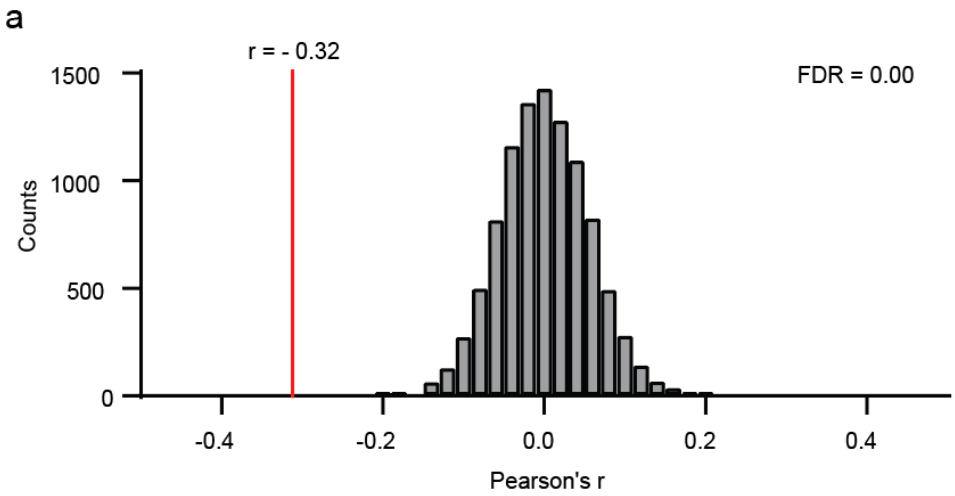


**Figure S2. The examination of the anti-correlation between intracellular strains.** (a) False discovery rate (FDR) was 0 using a Monte Carlo simulation that randomly permutes the value of nuclear strain and cytoplasmic strain for 300 cells, indicating that the anti-correlation between intracellular strains was not a false discovery.

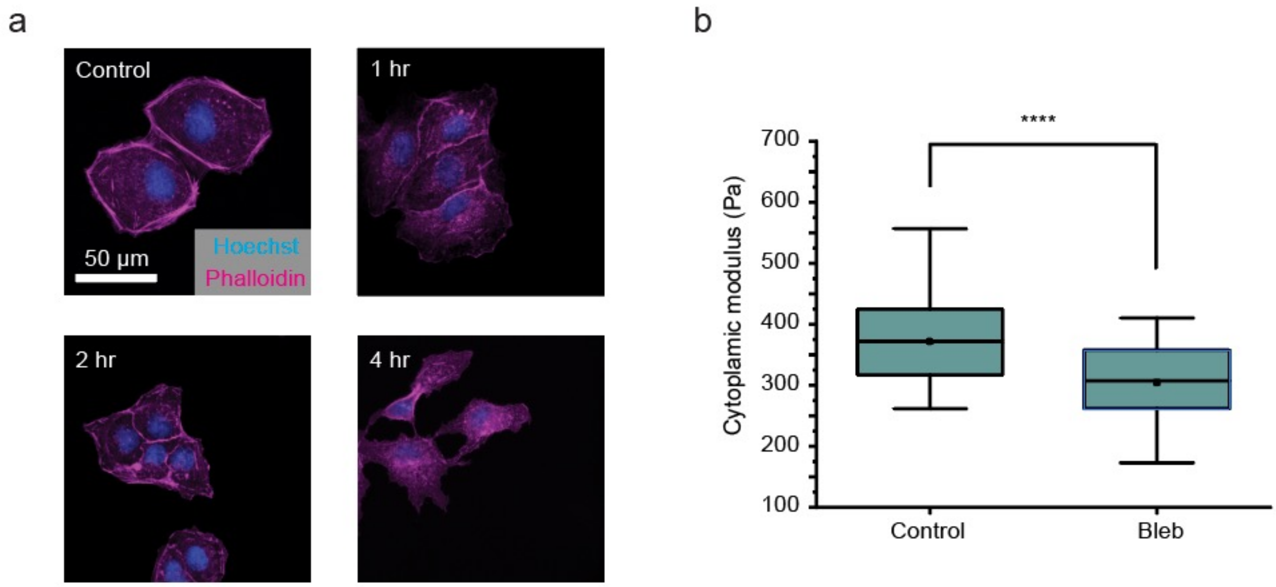


**Figure S3. The validation of blebbistatin treatment.** (a) Fluorescent images of actin filaments disrupted by 1, 2 and 4 hours blebbistatin treatments on MDCK cells. 2 hours treatment showed the best actin interference result and didn't significantly impact cell morphology as 4 hours treatment. (b) Cytoplasmic modulus was significantly reduced by blebbistatin treatment. For control and bleb, moduli were measured by AFM for N = 24 and N = 35 cells respectively.

a

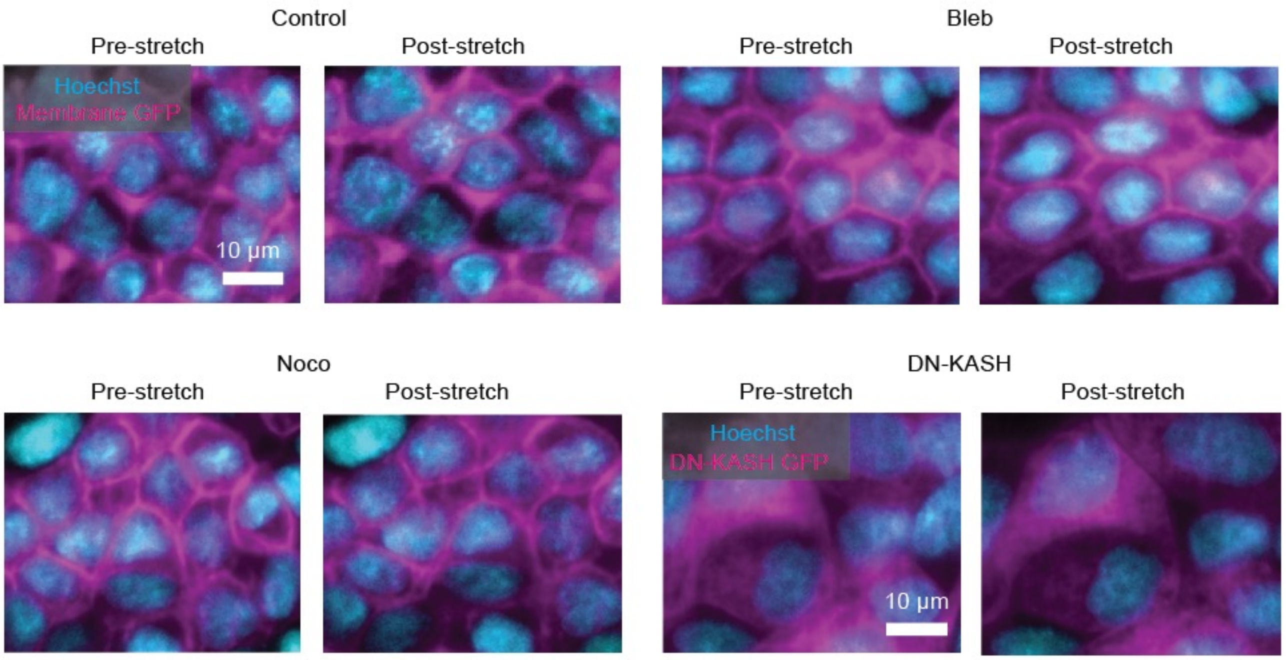


**Figure S4. Disruptions of intracellular force propagation** (a) Bleb, Noco, and DN-KASH showed no significant impact on cytoplasmic strain. (b) The representative images of the pre-stretch and post-stretch monolayers with or without the disruptions.

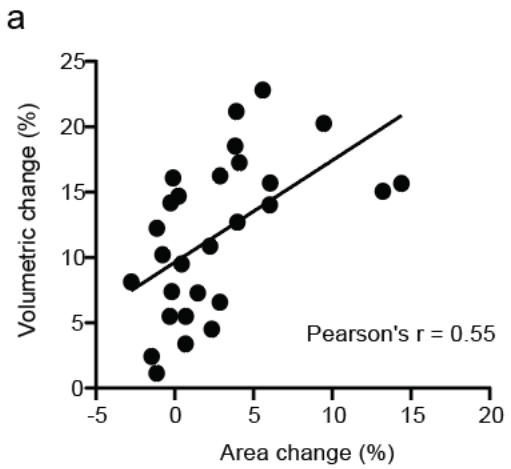


**Figure S5. Validation of the proportional correlation between nuclear area change and nuclear volumetric change.** (a) The scatter plot of nuclear area change versus nuclear volumetric change showed a high proportional correlation that the area change could be used for assessing the volumetric change.

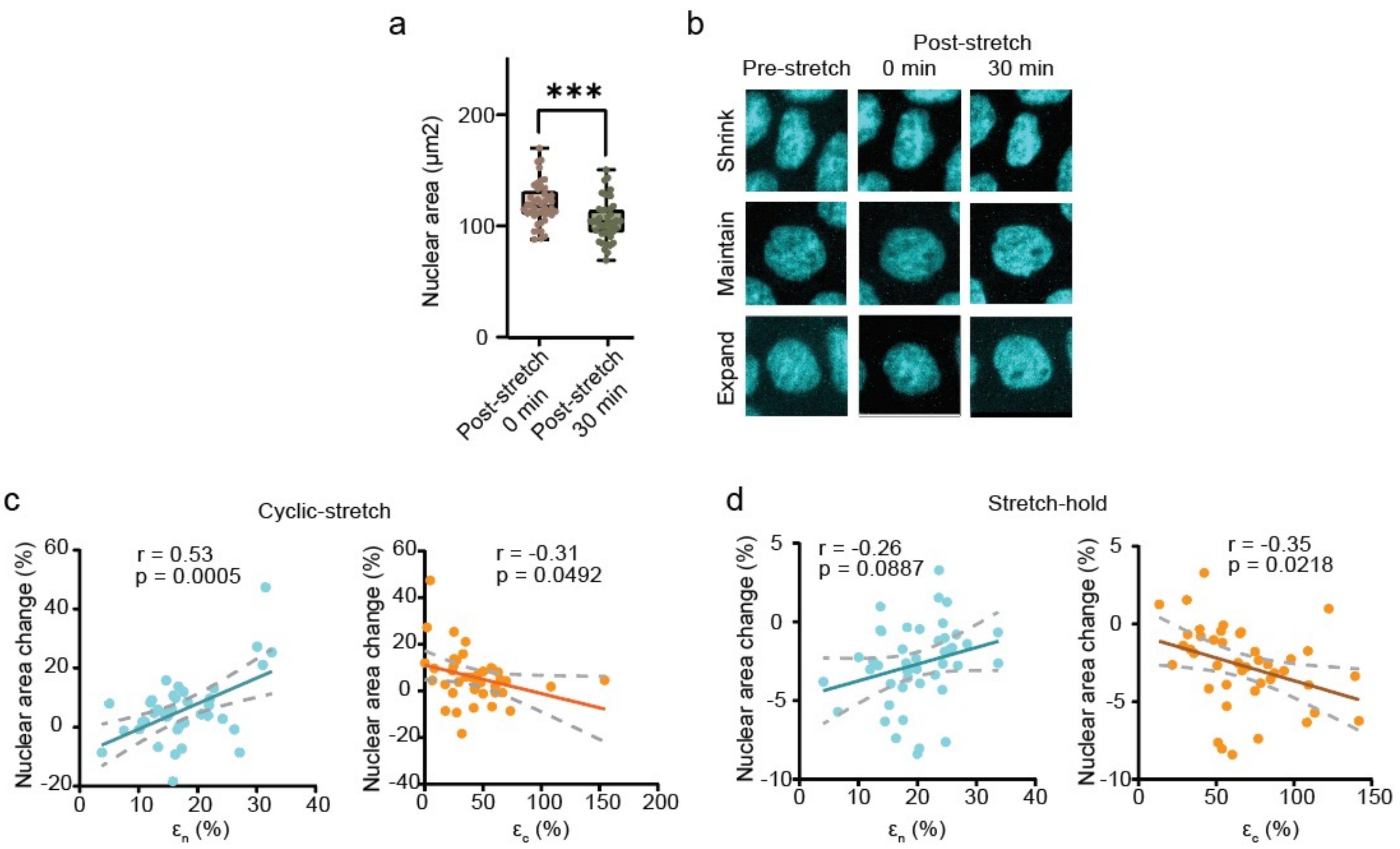


**Figure S6. Nuclei tended to shrink in stretch-hold experiment and the replication results.** (a) Nuclear area was reduced when holding the stretch over 30 min. N > 30 cells (b) Representative images of nuclear size change in stretch-hold experiment showed that nuclear could expand, shrink or maintain, similar to the observation in cyclic-stretch experiment. Replication of (c) cyclic-stretch experiment and (d) stretch-hold experiment showed the consistent results of the relationship between nuclear area change and intracellular strains.

| Features | Nucleus | Cell | Descriptions and Calculations |
|---|---|---|---|
| Perimeter | 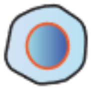 | 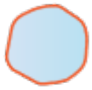 | The length of a closed path that outlines an object. |
| Area | 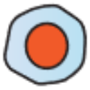 | 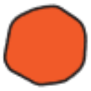 | The space taken up by the shape of an object. |
| Solidity | 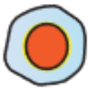 | 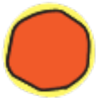 | The ratio of an object area to its convex hull. $Solidity = \frac{Area}{Convex\ hull}$. |
| Shape index | 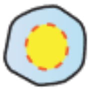 | 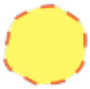 | The value indicates the shape of an object. $Shape\ index = \frac{Perimeter}{2 \times \sqrt{\pi \times Area}}$. |
| Circularity | 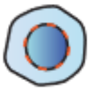 | 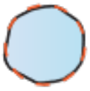 | The measurement of how closely the shape of an object approaches a circle. $Circularity = 4\,\pi \times \frac{Area}{Perimeter^2}$. |
| Aspect ratio | 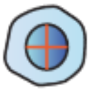 | 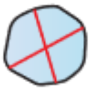 | The ratio of the major axis to minor axis in an object. |
| Roundness | 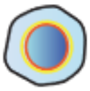 | 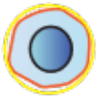 | The measurement of how closely an object approaches a circle with a diameter of the major axis. $Roundness = 4 \times \frac{Area}{\pi \times Major\ Axis^2}$. |
| Convex hull | 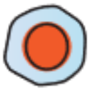 | 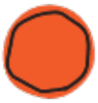 | The intersection of all convex sets that enclose an object. |
| Convex perimeter | 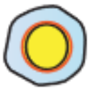 | 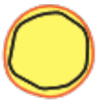 | The perimeter of a convex hull. |
| Nucleus-to-Cytoplasm area ratio | 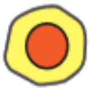 | | The ratio of the nucleus area to cytoplasm area. |

**Table S1. The morphological features in CCA.**
